# Communication through coherence by means of cross-frequency coupling

**DOI:** 10.1101/2020.03.09.984203

**Authors:** Joaquín González, Matias Cavelli, Alejandra Mondino, Nicolás Rubido, Adriano BL Tort, Pablo Torterolo

## Abstract

The theory of communication through coherence (CTC) posits the synchronization of brain oscillations as a key mechanism for information sharing and perceptual binding. In a parallel literature, hippocampal theta activity (4 – 10 Hz) has been shown to modulate the appearance of neocortical fast gamma oscillations (100 – 150 Hz), a phenomenon known as cross-frequency coupling (CFC). Even though CFC has also been previously associated with information routing, it remains to be determined whether it directly relates to CTC. In particular, for the theta-fast gamma example at hand, a critical question is to know if the phase of the theta cycle influences gamma synchronization across the neocortex. To answer this question, we designed a new screening method for detecting the modulation of the cross-regional high-frequency synchronization by the phase of slower oscillations. Upon applying the method, we found that the long-distance synchronization of neocortical fast gamma during REM sleep depends on the instantaneous phase of the theta rhythm. These results show that CFC is likely to aid long-range information transfer by facilitating the cross-regional synchronization of faster rhythms, thus consistent with classical CTC views.

## Introduction

Neuronal oscillations reflect changes in intracellular membrane potential excitability, and, as such, they provide windows of opportunity for efficient and energetically cheap information transmission (Buzsáki, 2006). Hence, long-distance communication in the brain is thought to depend on the synchronization of distributed neural networks at specific frequencies (Bastos et al., 2015; Engel et al., 2001). A possible mechanism is the so-called communication-through-coherence (CTC), which posits that phase coherence would be the neural substrate to achieve inter-regional information exchange (Fries, 2015, 2005). However, while the CTC only considers the coupling between oscillations of the same frequency, more recent work has categorically shown that oscillations of different frequencies also interact, a phenomenon called cross-frequency coupling (CFC) (Hyafil et al., 2015; Scheffer-Teixeira and Tort, 2018). For instance, the phase of the hippocampal theta rhythm (4–10 Hz) modulates the appearance of faster, gamma-frequency oscillations (30 – 150 Hz) through a process known as phase-amplitude coupling (Cavelli et al., 2018; Scheffer-Teixeira and Tort, 2017; Scheffzük et al., 2011; Tort et al., 2013).

In similarity to CTC, phase-amplitude CFC has also been linked information exchange in the brain (Canolty and Knight, 2010). But whether CTC and CFC would constitute independent or related processes remains to be better understood. It is reasonable to assume that if the coupling between brain rhythms of different frequencies aids long-range communication, it should also influence inter-regional phase coherence. To address this possibility, in the present work we designed a signal analysis metric to assess if the phase synchrony of fast oscillations (i.e., gamma) recorded in different regions is modulated by the phase of slower rhythms (i.e., theta). By applying this tool, we find that the phase of theta waves influences fast gamma (100 – 150 Hz, also called high-frequency oscillations) synchronization across the neocortex during REM sleep. These results suggest that the phenomena of CTC and CFC may be part of the same mechanism for efficient long-range communication.

## Results

To investigate the relation between CFC and CTC, we combined two well-established methods - the Modulation Index (MI) (Tort et al., 2010) and the Phase-Locking Value (PLV) (Lachaux et al., 1999; Varela et al., 2001) - into a new screening method to detect whether the phase of slower-frequency oscillations modulates long-range synchronization at higher frequencies. We employed this method on a dataset of 10 rats intracranially recorded during REM sleep. For simplicity, we focused on local field potential (LFP) recordings from three neocortical locations: the right V2 and the left and right S1 areas (Figure 1A). Standard spectral analyses (power, inter-regional synchrony, and phase-amplitude coupling) showed that these areas exhibit activity at theta and fast-gamma frequencies during REM sleep (Figure S1), as previously reported (Cavelli et al., 2018; Scheffer-Teixeira and Tort, 2017; Scheffzük et al., 2011).

**Figure 1.**
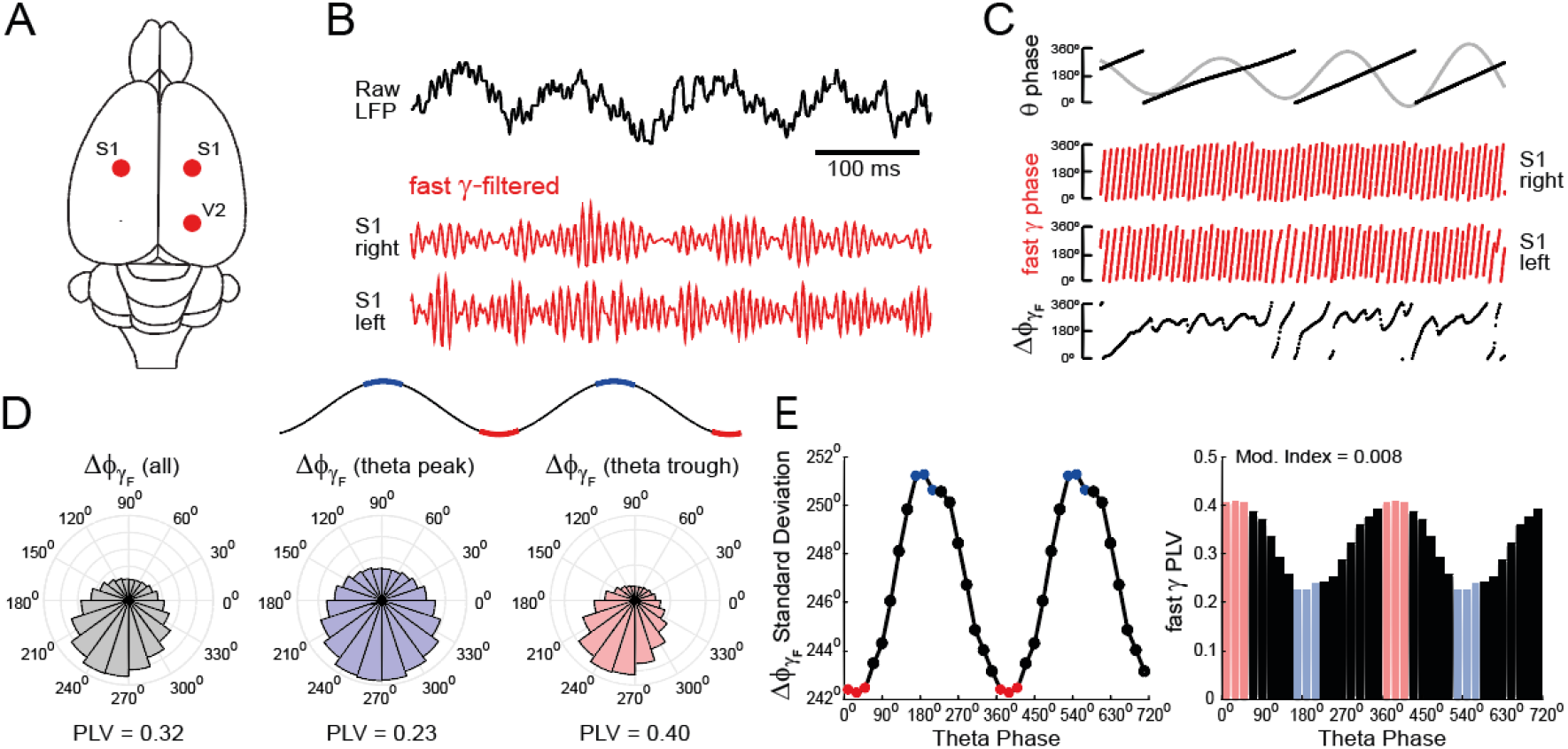
Detecting modulation of inter-regional gamma-frequency synchrony by the phase of theta. (A) Location of the neocortical electrodes analyzed in this work (S1, primary somatosensory cortex; V2, secondary visual cortex). (B) The top black trace shows an example theta rhythm (~8 Hz) readily observed during REM sleep in any raw neocortical LFP. The bottom red traces show the filtered fast-gamma (120-130 Hz) components from the left and right S1 areas. (C) The top panel shows the instantaneous phase (black) of the filtered theta oscillation (gray); the middle panels show the phases of the fast-gamma oscillations (red). The instantaneous inter-regional phase difference between the fast gamma oscillations (Δϕ_γ_) is shown at the bottom panel (black). Same time calibration as in B. (D) Circular distribution of Δϕ_γ_ for the entire REM sleep epoch (left, gray), as well as for when only analyzing Δϕ_γ_ values at the theta peaks (middle, blue) or troughs (right, red). The phase-locking value (PLV) obtained from each distribution is shown below. Notice the highest inter-regional gamma synchrony during the theta trough. (E) Left, Δϕ_γ_ standard deviation as a function of the theta phase. Notice the largest standard deviation near the theta wave peaks (lower synchronization) and the lowest near the trough (higher synchronization). Right, inter-regional fast-gamma PLV as a function of the theta phase. Colors depict the same phase bins employed in D (theta peak: blue; trough: red). The PLV modulation by the theta phase is quantified by means of a Modulation Index (see Methods).

Figure 1 outlines the new analysis framework, which is as follows: first, the LFPs from two distant brain regions are filtered to extract the high-frequency components (in the depicted example, we obtained the fast gamma activity from the left and right S1; Figure 1B). Then, the phase time series from these filtered signals are estimated by standard procedures (i.e., Hilbert transform; Figure 1C middle panels), which subsequently allows for the computation of their instantaneous phase difference (Δϕ_γ_; Figure 1C bottom panel). In classical approaches, the Δϕ_γ_ distribution is used to assess inter-regional synchrony through the PLV metric (Figure 1D left panel), which measures the concentration around the average Δϕ_γ_ value (Lachaux et al., 1999; Varela et al., 2001). Namely, the operational definition of synchrony at a given frequency is that of a constant phase difference between the regions. On the other hand, the less the two gamma activities are phase-locked, by definition the more uniformly distributed (over the circle) the Δϕ_γ_ values are, which renders its distribution less concentrated around the average phase difference.

The novelty introduced is the following one: instead of computing a single PLV using the entire Δϕ_γ_ time series, we computed several PLVs using non-overlapping subsets of Δϕ_γ_ values. Note in the bottom panel of Figure 1C that the instantaneous Δϕ_γ_ values fluctuate in time between periods of roughly constant values (high synchronization) and those of phase advancements and delays (low synchronization). We sought to determine whether such fluctuations would be related to the theta cycle. To that end, we obtained the phase time series of a reference LFP signal filtered at theta frequency (Figure 1C top; we used the S1 LFP as the reference signal, which has the same theta phase as the hippocampus stratum oriens). Next, we computed Δϕ_γ_ distributions for different phases of the theta cycle (Figure 1D right panels). Notice that, for this example REM sleep epoch, the inter-regional fast-gamma synchrony substantially changes across the theta cycle. For instance, Δϕ_γ_ concentration is almost twice near the theta through than at the theta peak (as assessed by the PLV; Figure 1D). Notice further in Figure 1D a less concentrated Δϕ_γ_ distribution near the theta peak (blue) with a resultant lower PLV compared to the average global PLV (gray), and the opposite happening for the theta trough (red). Of note, the preferred phase difference (i.e., the average Δϕ_γ_) does not significantly differ across the theta cycle (for Figure 1D example: global Δϕ_γ_ average: 257°; average Δϕ_γ_ at the peak: 267°; trough: 252°; at the group level [n=10 rats]: global Δϕ_γ_ average: 195° ± 15°; peak: 190° ± 22°; trough: 196° ± 11°; see Figure S2).

Figure 1E depicts the circular standard deviation of Δϕ_γ_ (left panel) as well as the PLV distribution as a function of the theta phase (right panel). Both panels show that the phase of the theta cycle modulates the inter-regional fast-gamma synchrony between the left and right S1 areas. To quantify this synchrony modulation, the last step constitutes applying the MI to the PLV-phase distribution. As in related approaches (Tort et al., 2010), a null MI implies a uniform PLV distribution; that is, the PLV does not depend on theta and has the same value across phases. On the other hand, the higher the PLV modulation, the higher the MI. The maximal possible MI value is 1, which is achieved for the case in which the PLV is different from zero only in a single phase bin and equals zero otherwise. However, similarly to phase-amplitude comodulation, such a case is rather theoretical and - for biological networks - the statistically significant MI values are usually in the order of 10^−3^ to 10^−2^(Scheffer-Teixeira and Tort, 2018).

We next repeated the procedure described above for nine different pairs of high- and low-frequency rhythms. The results revealed that only the phase of theta (and not delta nor alpha) influenced long-range synchronization during REM sleep; moreover, such synchrony modulation was specific for the fast-gamma activity (Figure 2A). In order to perform a group level analysis (n=10 rats), we computed phase-synchrony comodulation maps (or “comodulograms”) for each individual animal, which were achieved by plotting the MI of several frequency pairs in pseudocolor scale. The significance of MI values of each comodulogram was then assessed by a surrogate analysis (see Methods and Figure S3). In Figure 2B, we plot the average comodulogram across animals; the transparent line demarks the phase-synchrony couplings that were statistically significant in all animals. Note coupling specificity for theta phase and fast-gamma synchrony. Interestingly, theta modulation was much higher for inter-hemispheric (S1-S1) than intra-hemispheric (S1-V2) long-range synchrony (p=0.002, Wilcoxon signed-rank test; Figure 2C).

**Figure 2.**
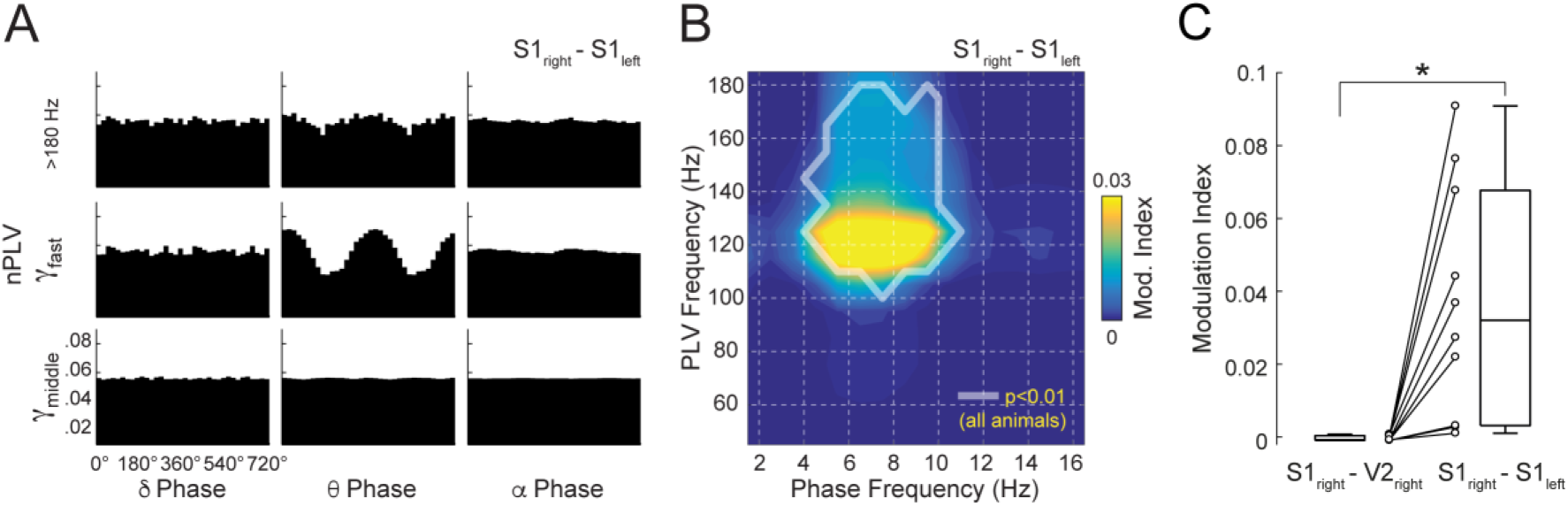
Theta waves specifically modulate neocortical fast-gamma synchronization during REM sleep. (A) Normalized phase-locking value (nPLV) distributions for three fast oscillatory sub-bands as a function of the phase of three slow frequencies. Each panel shows a fast-slow rhythm combination; 9 combinations are shown which include middle gamma (60-80 Hz), fast gamma (120-130 Hz), and oscillations >180 Hz combined with the delta (1-4 Hz), theta (4-10 Hz) and alpha (10-15 Hz) frequency bands. For each fast-slow frequency pair, the PLV normalization is obtained by dividing individual phase bin values by the sum across bins. Notice that inter-regional synchrony modulation occurs mostly between fast-gamma activity and theta phase. Results correspond to the analysis of the right and left S1 electrodes of a representative animal during REM sleep. (B) Population average (n = 10 rats) phase-synchrony comodulation map computed for inter-hemispheric S1 synchrony during REM sleep. The x-axis represents the reference phase frequency and the y-axis the fast frequency analyzed for synchrony through the PLV. The level of phase synchrony modulation is quantified by the MI and shown in pseudocolor code. The transparent polygon circumscribes frequency pairs that were statistically significant (p<0.01) in all animals. Significance thresholds were obtained through a surrogate analysis (see Methods and Figure S3 for details). (C) Modulation index of theta-fast gamma synchronization for intra- and inter-hemispheric locations. Theta modulation is much higher for inter-hemispheric gamma synchrony (*p=0.002, Wilcoxon signed-rank test, n=10 rats).

## Discussion

In the present work, we sought to investigate the functional relationship between CFC and CTC, two network mechanisms that have been independently hypothesized to underlie information exchange (Canolty and Knight, 2010; Fries, 2005). We developed a new signal analysis metric to detect the modulation of cross-regional synchronization of fast oscillations by the phase of slower rhythms. Our results reveal that the previously separately described CTC and CFC phenomena are related. In specific, we found that CFC links not only slow and fast oscillations but also inter-regional coherence at fast frequencies. In this regard, we observed that slow-wave cycles generate transient periods of enhanced long-range synchronization at higher frequencies, which is to say that the phase of slow rhythms determines the temporal windows for efficient CTC to take place.

To illustrate the usefulness of our method, we examined the well-known case of theta-fast gamma coupling during REM sleep, in which the phase of theta modulates the amplitude of neocortical 110-150 Hz oscillations (Cavelli et al., 2018; Scheffer-Teixeira and Tort, 2017; Scheffzük et al., 2011; Tort et al., 2013). Our novel tool expands this consolidated result by showing that the theta phase also modulates inter-regional phase coherence at fast gamma frequency during REM sleep (Figure 1). Such an effect is frequency-specific since it did not occur for other slow-fast frequency pairs, nor other gamma sub-bands (Figure 2). Moreover, we found that theta oscillations have a stronger influence over inter-hemispheric than intra-hemispheric synchrony (Figure S4), suggesting that theta may be particularly well suited for connecting information processed in both hemispheres.

In rodents, theta oscillations occur during active wakefulness and REM sleep states, and have been linked to spatial navigation and memory consolidation (Buzsáki and Moser, 2013). At the neuronal level, theta oscillations orchestrate firing patterns and entrain gamma rhythms in the hippocampus and neocortex (Scheffer-Teixeira et al., 2012; Schomburg et al., 2014; Sirota et al., 2008). Our results add to these observations by showing that theta also modulates inter-regional gamma synchrony during REM sleep. Such an effect may facilitate neocortical information routing during memory consolidation, in line with reports pointing to the importance of theta to memory consolidation during sleep (Boyce et al., 2016; Nishida et al., 2008). We speculate that theta-modulated CTC may help to consolidate memories stored
 through CFC patterns (Axmacher et al., 2010; Lega et al., 2014; Lisman and Jensen, 2013; Tort et al., 2009, 2008; van Wingerden et al., 2014) into the neocortex as distributed cell assemblies (Benchenane et al., 2010; Canolty et al., 2010).

Related evidence has been previously observed in scalp EEG recordings of human subjects by Doesburg and colleagues. These authors showed that the phase of theta range activity (~6 Hz) modulates inter-regional EEG synchronization at the low gamma band (30-50 Hz) when humans perform auditory, visual and verbal tasks (Doesburg et al., 2012a, 2009; see also Doesburg et al., 2012b). Compared to the prominent effects found here in rats, however, the statistical significance of their synchrony modulation was rather marginal (e.g., see Figure 7 in Doesburg et al., 2009 and Figure 6 in Doesburg et al., 2012a). Moreover, it is unclear whether the EEG recordings exhibited actual power peaks at theta and gamma (c.f. Figure S1) so as to confidently infer the presence of genuine oscillatory activity (see Yuval-Greenberg et al., 2008), nor whether there was any frequency specificity for the analyzed bands. In addition to these findings in humans, Bosman et al. (2012) provided preliminary evidence from two macaque monkeys showing that V1-V4 coherence at 70-80 Hz depends on the time from the peaks of the 3-5 Hz-filtered LFP component. Furthermore, we have also recently shown that the phase of respiratory activity modulates low gamma synchrony in the awake cat (Cavelli et al., 2019). Altogether, the current and previous evidence suggests that the rhythmic modulation of phase synchrony may be a general brain mechanism preserved across species.

As a cautionary remark, our phase-synchrony metric detects PLV variations within the cycles of slower oscillations, but it is independent of their absolute values. This means that a statistically significant synchrony modulation may be detected even in the presence of low synchrony levels. In fact, we showed that inter-hemispheric fast-gamma coherence has higher theta modulation than the intra-hemispheric coherence (Figure 2C), despite the absolute synchrony level being higher for the latter (Figure S1). Finally, we also remark that a phase-modulated PLV does not imply that phase-locking occurs only at a particular phase, but that varying levels of phase-locking can be observed across the slow oscillation cycle.

In summary, here we described a new method that can be generally applied to any brain rhythms as a screening tool for finding whether the phase of slower oscillations influences phase synchronization at faster frequencies. This work opens the possibility to revisit former CFC results under the prism of their functional relationship with CTC.

## Acknowledgments

This study was supported by the Programa de Desarrollo de Ciencias Básicas, PEDECIBA; Agencia Nacional de Investigación e Innovación (ANII), (FCE_1_2017_1_136550) and the Comisión Sectorial de Investigación Científica (CSIC) I+D-2016-589 grant from Uruguay. J.G was supported by CAP (Comisión Académica de Posgrado). N.R. acknowledges the CSIC group grant CSIC2018 - FID 13 - Grupo ID 722. A.B.L.T. was supported by Conselho Nacional de Desenvolvimento Cientifico e Tecnológico (CNPq) and Coordenação de Aperfeiçoamento de Pessoal de Nível Superior (CAPES), Brazil.

## Author Contributions

J.G., M.C and P.T. designed the experiments; J.G., M.C. and A.M conducted the experiments; J.G. and A.B.L.T. wrote analysis software; J.G. analyzed the data; J.G., M.C., N.R., A.B.L.T. and P.T. were involved in the discussion and interpretation of the results; J.G, A.B.L.T. and P.T. wrote the manuscript.

## Code and Data availability

Matlab codes for the performing the analysis framework described here are freely available at https://github.com/tortlab/Phase-Locking-Value---Modulation-Index. Data is available upon reasonable request to the authors.

### Methods

#### Experimental animals

We used 10 male Wistar adult rats, maintained on a 12-h light/dark cycle (lights on at 07:00 am) with food and water freely available. We conducted all experimental procedures in agreement with the National Animal Care Law (No. 18611) and with the *Guide to the care and use of laboratory animals* (8^th^ edition, National Academy Press, Washington DC, 2010). Furthermore, the Institutional Animal Care Committee (Comisión de Etica en el Uso de Animales) approved the experiments (Exp. No 070153-000332-16). The animals were determined to be in good health by veterinarians of the institution. We also took all available measures to minimize pain, discomfort, and stress, while employing the minimum number of animals necessary to obtain reliable scientific data.

#### Surgical Procedures

We implanted the animals with intracranial electrodes through similar surgical procedures as in previous studies (Cavelli et al., 2018). The anesthetic was a mixture of ketamine-xylazine (90 mg/kg and 5 mg/kg i.p., respectively). We positioned the rat in a stereotaxic frame, exposed the skull and placed stainless steel screws directly above somatosensory (S1) and visual (V2) cortices (bilateral); S1 and V2 coordinates from bregma were −2.5 mm and −7.5 mm anteroposterior, respectively; all electrodes were placed ±2.5 mm from the midline. Additionally, we placed another steel screw above the cerebellum to work as a reference, and two electrodes into the neck muscle to record the electromyogram (EMG). All electrodes were soldered into a 12-pin socket and fixed onto the skull with acrylic cement. At the end of the surgical procedures, we administered Ketoprofen (1 mg/kg s.c.) to reduce pain. After the animals had recovered from surgery, they were adapted to the recording chamber for one week.

#### Experimental Sessions

For the recordings, the animals were placed inside transparent cages (40×30× 20 cm) containing wood shaving material in a temperature-controlled (21-24°C) room, with water and food *ad libitum*. Each experimental session took place during the light period, between 10 am and 4 pm in a sound-attenuated chamber with a Faraday shield. During the experiments, the animals were connected through a rotating connector which allowed them to move freely. The recordings were amplified (x1000), acquired and stored in a computer using Dasy Lab Software through a 16-bit AD converter (sampling frequency: 1024 Hz).

#### Sleep staging

We classified the states of sleep and wakefulness by analyzing electrophysiological and EMG recordings in 10-s epochs. We followed the same criteria as in our previous work (Cavelli et al., 2018). In brief, wakefulness was characterized by relatively high EMG activity; NREM by high-voltage slow cortical waves together with sleep spindles and reduced EMG amplitude; REM sleep was characterized by regular theta activity and silent EMG, except for occasional twitches. After this first staging, we subsequently discarded epochs exhibiting recording artifacts. For each animal, we concatenated all artifact-free REM sleep epochs before carrying the analysis described below.

#### Data analysis

We developed a metric to detect whether the phase of a slow oscillation modulates the phase-synchronization of a higher frequency rhythm between two areas. Our metric was constructed by combining two previous methods: the Modulation Index (MI, Tort et al., 2010) and the Phase-Locking Value (PLV, Lachaux et al., 1999).

As is the case of the original MI and PLV metrics, the new method consists in analyzing filtered signals. In this work, low- and high-frequency LFP components were obtained by band-pass filtering using the eegfilt() Matlab function (Delorme and Makeig, 2004) with 3-Hz and 10-Hz bandwidths, respectively. The phase time series of the filtered signals were then estimated from their analytical representation based on the Hilbert transform. In the new framework, three phase time series need to be estimated: one for the low-frequency signal (e.g., a theta reference signal) and one for each of the two high-frequency signals (e.g., gamma activity from two different brain regions). Note that this step contrasts with the original MI formulation, in which the high-frequency amplitudes and not the phases are employed (Tort et al., 2010).

After estimating the phase time series, we computed the PLV between the high-frequency components as a function of the phase of the low-frequency component. We accomplished this by partitioning the phase of the slower oscillation into 18 bins and computing PLVs using subsets of the high-frequency phase time series restricted to each of the phase bins. Similar results are obtained when employing a different number of phase bins (see Figure S4).

The PLV for the i-th bin is defined as:

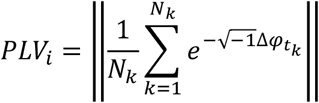

Where 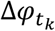 is the instantaneous phase difference between the two fast oscillations at time 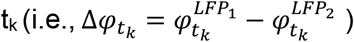. The summation is carried across all time stamps t_k_ associated with the phase bin i, and N_k_ is the total number of data points.

We next applied a normalization by dividing the PLV of each phase bin by the sum across all phase bins:

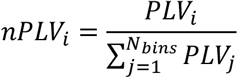

Where N_bins_ is the number of phase bins (in our case, N_bins_=18). We then used the MI to quantify the PLV-phase modulation (“MI_PLV_”). This was achieved by first computing the Shannon Entropy (H) of the nPLV phase-distribution:

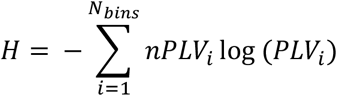

And then:

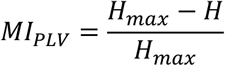

Where H_max_ = log(N_bins_) is the maximum entropy, which corresponds to a uniform nPLV distribution and no phase modulation. Hence, the MI_PLV_ measures how far is the nPLV distribution from a uniform one.

Finally, to obtain the phase-synchrony comodulogram, we repeated the procedure above for several frequency pairs and expressed the MI_PLV_ values by means of a pseudocolor scale. Low-frequency components were obtained in steps of 1 Hz, from 1 to 16 Hz, and high-frequency components in 10 Hz steps, from 40 to 180 Hz.

#### Statistical analysis

To detect the statistical significance of MI_PLV_ values, we performed a surrogate analysis to obtain a null hypothesis distribution (see Figure S3). Surrogates were constructed by randomly circularly shifting the slow oscillation phase between 1 and 10 seconds before computing the MI_PLV_. This procedure was repeated 100 times per animal, and, for each animal, we set p<0.01 as our statistical threshold, meaning that the original MI_PLV_ value had to be higher than the 100 surrogate values. The levels of inter- and intra-hemispheric synchrony modulation were compared by means of the Wilcoxon signed-rank test.

## Supplementary Material

**Figure S1.**
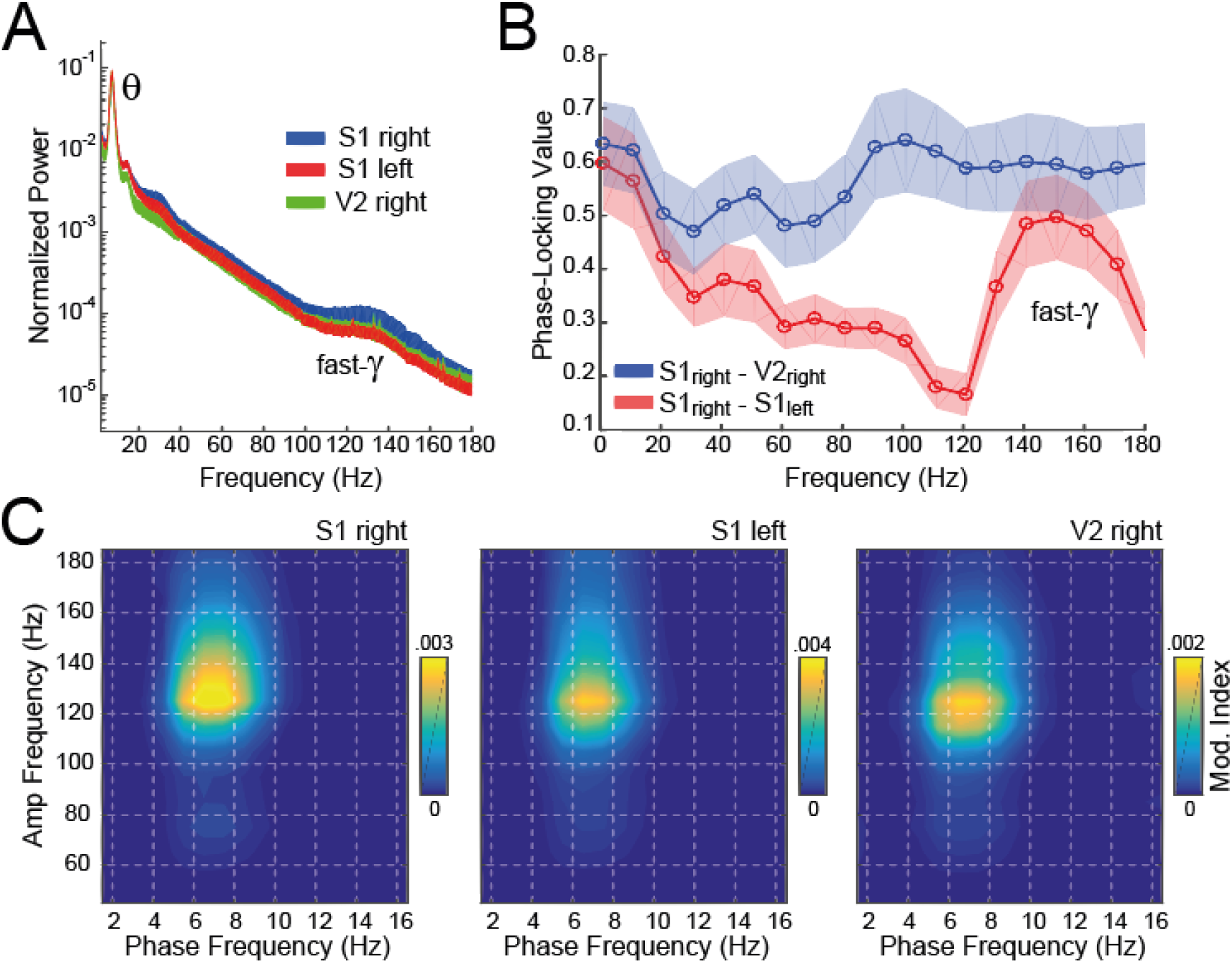
Standard LFP analysis during REM sleep. (A) Normalized power spectra (mean ± SEM, n=10 rats). Notice power peaks at theta and fast-gamma in all neocortical locations. The normalization was performed within each animal by dividing power values by the sum across frequencies. (B) Average (± SEM) phase-locking value (PLV) as a function of frequency. PLV was calculated for the intra-hemispheric (S1-V2, blue) and the inter-hemispheric (S1-S1, red) electrode combinations. Circles denote the filter center frequency (bandwidth: 10 Hz). (C) Average phase-amplitude comodulograms for REM sleep epochs (computed as described in Tort et al., 2010). Note the theta modulation of fast-gamma amplitude in all analyzed neocortical regions, as previously reported (Cavelli et al., 2018; Scheffer-Teixeira and Tort, 2017; Scheffzük et al., 2011).

**Figure S2.**
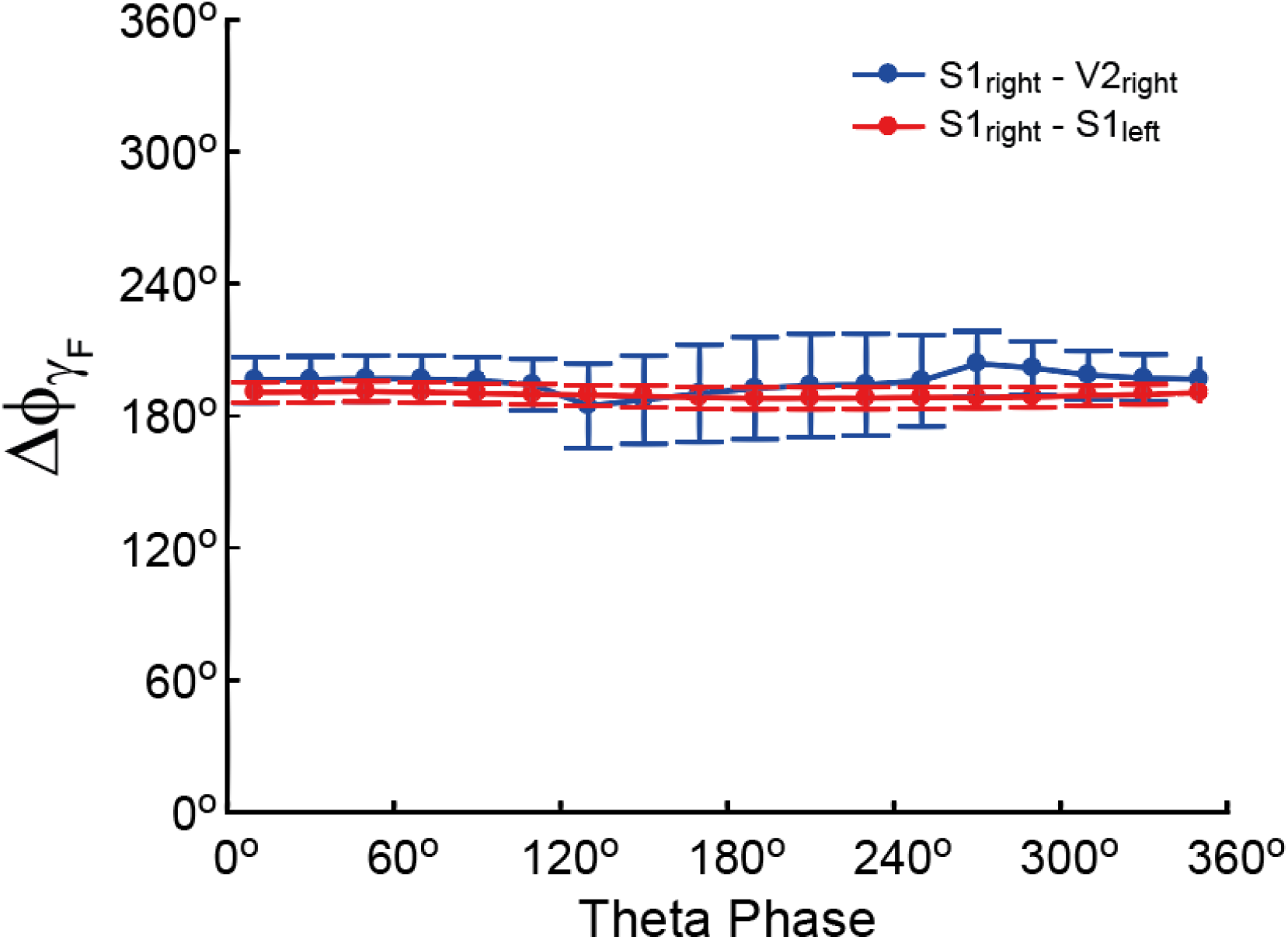
Similar gamma phase difference across the theta cycle. Average phase difference (Δϕ_γ_) as a function of theta phase for fast gamma oscillations recorded from two distant electrodes located in the same (blue) and in different hemispheres (red).

**Figure S3.**
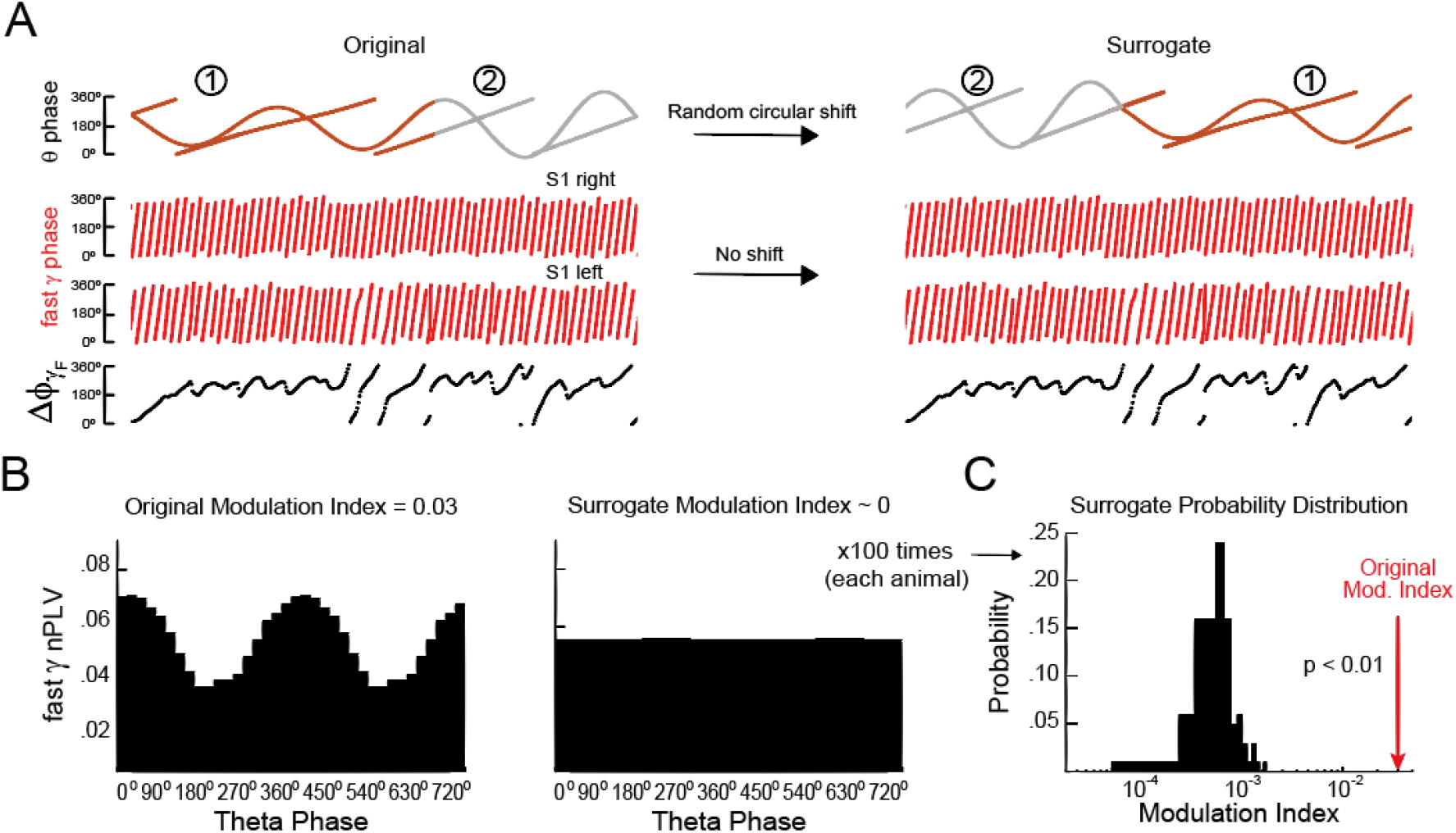
Schematics of surrogate analysis for statistical testing of modulation index (MI) values. (A) To test MI significance, we employed a surrogate analysis based on circularly shifting the slow oscillation phase while keeping the phase of the fast oscillations unaltered. (B) Normalized PLV distribution over the theta phase computed for the original and surrogate time series. Notice lack of phase-synchrony modulation in the latter. (C) For each analyzed pair of slow and fast frequencies, the circular shift was performed 100 times per animal, leading to a distribution of 100 surrogate MI values. The original MI is then compared to the surrogate distribution. We assume p<0.01 when the original MI value is larger than all 100 surrogate values.

**Figure S4.**
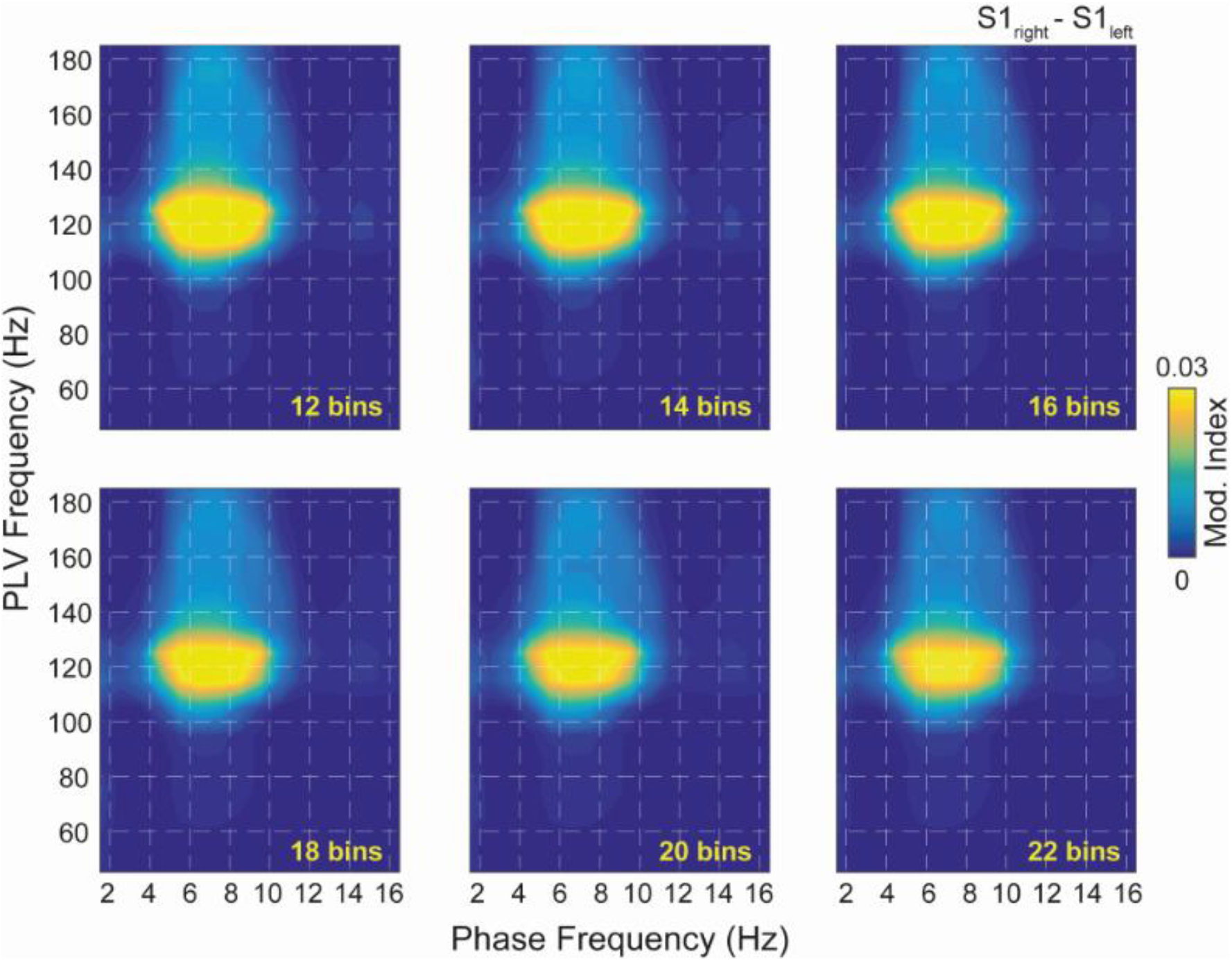
Similar phase-synchrony comodulograms for different theta cycle partitions. Each comodulogram was obtained by dividing the theta cycle into a different number of phase bins, from 12 to 22 bins (bin length range: 30° to 16°).

